# Highly sensitive quantitative phase microscopy and deep learning complement whole genome sequencing for rapid detection of infection and antimicrobial resistance

**DOI:** 10.1101/2022.07.07.499154

**Authors:** Azeem Ahmad, Ramith Hettiarachchi, Abdolrahman Khezri, Balpreet Singh Ahluwalia, Dushan N. Wadduwage, Rafi Ahmad

**Affiliations:** Department of Physics and Technology, UiT The Arctic University of Norway, Tromsø, 9037, Norway; Department of Electronic and Telecommunication Engineering, University of Moratuwa, Sri Lanka; Department of Biotechnology, Inland Norway University of Applied Sciences, Holsetgata 22, 2317, Hamar, Norway; Department of Clinical Science, Intervention and Technology, Karolinska Insitute, 17177 Stockholm, Sweden; Center for Advanced Imaging, Faculty of Arts and Sciences, Harvard University, Cambridge, USA; Institute of Clinical Medicine, Faculty of Health Sciences, UiT - The Arctic University of Norway, 9037, Norway

**Author notes:** These authors contributed equally (Ramith Hettiarachchi, Abdolrahman Khezri). These authors contributed equally (Balpreet Singh Ahluwalia, Dushan N. Wadduwage, Rafi Ahmad).

## Abstract

The current state-of-the-art infection and antimicrobial resistance diagnostics (AMR) is based mainly on culture-based methods with a detection time of 48-96 hours. Slow diagnoses lead to adverse patient outcomes that directly correlate with the time taken to administer optimal antimicrobials. Mortality risk doubles with a 24-hour delay in providing appropriate antibiotics in cases of bacteremia. Therefore, it is essential to develop novel methods that can promptly and accurately diagnose microbial infections at both species and strain levels in clinical settings. Here, we demonstrate that the complimentary use of label-free optical assay with whole-genome sequencing (WGS) can enable high-speed culture-free diagnosis of infection and AMR. Our assay is based on microscopy methods exploiting label-free, highly sensitive quantitative phase microscopy (QPM) followed by deep convolutional neural networks (DCNNs) based classification. We benchmarked our proposed workflow on 21 clinical isolates from four WHO priority pathogens (*Escherichia coli, Staphylococcus aureus, Klebsiella pneumoniae*, and *Acinetobacter baumannii*) that were antibiotic susceptibility testing (AST) phenotyped, and their antimicrobial resistance (AMR) profile was determined by WGS. The proposed optical assay was in good agreement with the WGS characterization. Highly accurate classification based on the gram staining (100% for gram-negative and 83.4% for gram-positive), species (98.6%), and resistant/susceptible type (96.4%), as well as at the individual strain level (100% accurate in predicting 19 out of the 21 strains). These results demonstrate the potential of the QPM assay as a rapid and first-stage tool for species, presence, and absence of AMR, and strain-level classification, which WGS can follow up for confirmation of the pathogen ID and the characterization of the AMR profile and susceptibility antibiotic. Taken together, all this information is of high clinical importance. Such a workflow could potentially facilitate efficient antimicrobial stewardship and prevent the spread of AMR.

## 1. Introduction

Antimicrobial resistance (AMR) is the ability of microorganisms to resist antimicrobial treatments, especially antibiotics. Infections due to antibiotic-resistant bacteria are a threat to modern healthcare. A recent meta-analysis of resistant bacteria burden on human health and well-being has revealed that in 2019 alone, 1.27 million deaths were caused directly by antibiotic-resistant bacteria (ARBs), and 4.95 million deaths were associated with ARBs [1]. This number has surpassed HIV and malaria. Many (73%) of these deaths are due to infections caused by *Escherichia coli, Staphylococcus aureus, Klebsiella pneumoniae, Acinetobacter baumannii*, and *Pseudomonas aeruginosa* [2], which the WHO identifies as critical and high-priority pathogen groups.

Research has shown that adverse patient outcome directly correlates with the time taken to administer optimal antimicrobial [3]. Mortality risk doubles with a 24-hour delay in providing appropriate antibiotics in cases of bacteremia [4]. Globally only half of the antibiotics are prescribed correctly [5]. Thus, rapid point-of-care diagnostic tests are a central part of the solution to this problem. Current culture-based methods used to detect and identify agents of infection are inadequate. Incubation times of up to 24-48 hours are often necessary to capture the majority of culturable bacteria associated with infection [6]. Additional time is required for pathogen identifications (2-4h) and in case of expected AMR for antibiotic susceptibility testing (AST) (18-24h). Thus, the time interval from collecting patient samples at the ward until the information is available on antibiotic susceptibility patterns is, in the best case, 2-4 days in the clinical routine [7].

There are emerging micro-and nanotechnologies for bacterial identification and AST, including both phenotypic (e.g., microfluidic-based bacterial culture) and molecular (e.g., multiplex PCR, hybridization probes, nanoparticles, synthetic biology, and mass spectrometry) methods [8]. PCR and MS methods used on positive cultures to identify microbes have progressed considerably but still are far from ideal. With PCR, one needs to decide beforehand what to look for, and MALDI-TOF is expensive [6, 9].

Whole genome sequencing (WGS) can overcome some of these problems since there is no need for targeted primers/probes. Also, with the rise of real-time sequencing and its affordability, WGS has become a potent alternative to time-consuming culture-dependent traditional methods. We have recently demonstrated that using Oxford Nanopore Technologies (ONT) MinION and Flongle sequencing platforms, infectious agents and their resistance profile can be identified within 10 min – 1 hour after the start of the sequencing [6, 10, 11]. Additionally, we have used data from direct sequencing of spiked blood cultures for genotype-to-phenotype prediction of resistance towards B-lactams in *E. coli* and *K. pneumoniae*, as fast as 1–8 h from the sequencing start. However, around 3-4 hours are still required to prepare the sample for sequencing.

Direct identification of the pathogen in biofluid samples will mitigate the above issues. Culture-independent diagnostic tests can also detect dead bacteria when antibiotic therapy has been administered before sampling, which could help detect the pathogen [12]. Real-time sequencing, for example, has been successfully implemented in lower respiratory tract infections, urine infections, cerebral spinal fluid, surgical site infections, and orthopedic devices with up to 100 % sensitivity and specificity in pathogen detection. Label-free optical techniques such as quantitative phase microscopy [13] and Raman spectroscopy [14, 15] have recently been shown to measure phenotypic and molecular signatures in pathogens at very low concentrations. We have recently reported the multi-excitation Raman spectroscopy (ME-RS) method for the species, resistance, and strain-level classification of pathogens [16]. In combination with downstream machine learning-based analysis, such methods may open doors to developing new culture-free identification of bacteria strains and their clinically relevant properties, such as wild-type (WT) and non-wild type (NWT) bacteria and provide insight into the AMR profile.

Quantitative phase microscopy (QPM) is a powerful technique due to its non-contact, non-invasive, label-free, and quantitative nature. This makes QPM a suitable candidate in the domain of biomedical imaging applications. QPM can measure various morphological (surface area, volume, sphericity, etc.), bio-physical (dry mass, growth rate, etc.), and statistical measures of morphological (mean, standard deviation, skewness, kurtosis, etc.) parameters related to cells/tissues at nanometric sensitivity [17-20]. QPM encodes information about the specimens’ optical thickness (refractive index × geometrical thickness) in terms of modulated intensity patterns called interferograms. There exist various optical configurations of QPM, such as diffraction phase microscopy [20], spatial light interference microscopy [21], Linnik interference microscopy [22-24], Mirau interference microscopy [25-27], and Mach-Zehnder interference microscopy [28].

The spatial phase sensitivity of the QPM system is the most crucial parameter which decides the minimal detectable change in the cell’s parameters either as a function of their growth or different categories such as normal and challenged. The phase sensitivity of QPM depends on the type of light source (white light, LEDs, and laser) used to illuminate the specimens for the generation of raw data, i.e., interferograms. These light sources either generate highly sensitive phase images at the cost of reduced temporal resolution (for white light and LEDs) or provide high temporal resolution at the cost of less phase sensitivity (for laser) [23]. The pros and cons of these light sources in QPM have been discussed in great detail in Refs. [23, 24]. Recently, a temporally high and spatially low coherent light source, also known as a pseudo-thermal light source (PTLS), has been proposed to resolve the issues associated with using the aforementioned light sources in QPM [23, 25, 26, 29-31]. PTLS can provide highly sensitive phase images comparable to white light illumination without sacrificing temporal resolution. Despite the apparent advantages of QPM, it is still challenging to differentiate the large number of classes of biological samples based on the manually quantified cell parameters mentioned in the previous paragraph. Therefore, the extracted phase images of different categories of the bio-specimens and WGS can be utilized for their classification by employing a powerful computational approach, e.g., machine learning or deep learning. With the larger amount of data generated from QPM and WGS, robust methods for data analysis and correlation must be needed to generate clinically relevant results.

Deep learning methods have revolutionized all scientific fields, including bio-imaging, genomics, and computer vision. Deep convolutional neural networks (DCNNs) outperformed all classical methods for vision tasks, such as image segmentation, semantic segmentation, and classification. DCNNs are also increasingly being adopted in microscopy, including in QPM [32]. Jo *et al*. proposed a DCNN to classify quantitative phase images [32]. Their results show that deep learning over conventional machine learning (ML) techniques can improve the classification performance between five species of Bacillus (*B. anthracis, B. thuringiensis, B. cereus, B. atrophaeus*, and *B. subtilis*). Although the network identified *B. anthracis* with high accuracy (∼80%), the accuracy ranged between 45% and 75% for other species. Kim *et al*. proposed a CNN-based framework that uses 3D refractive index tomograms to identify 19 types of bacteria and properties such as gram staining and aerobicity [33]. The authors observed less confusion between bacteria species of the same genus. In a study by Wang *et al*., they designed a computational live bacteria detection system that periodically captures microscopy images of bacterial growth inside an agar plate [34]. Within 12 h of incubation, their classification network has obtained species-specific accuracies of ∼97.2%, ∼84.0%, and ∼98.5% for *E. coli, K. aerogenes*, and *K. pneumoniae*, respectively.

This study developed a label-free optical assay to identify bacteria and their clinically relevant culture-free properties. Our assay is based on partially spatially coherent high-sensitive QPM followed by DCNN-based classification. First, various challenges associated with the bacteria phase imaging are resolved, such as the effect of defocusing and background aberration removal for error-free phase map generation. The QPM experiments are then performed on 21 types of bacteria strains, which had been previously genotype and phenotype. Our results suggest that the proposed assay can classify 19 of the 21 strains with 100% accuracy at extremely low bacteria concentrations (less than 100 bacteria per sample). For the authenticity of the classification process, we further correlated the interclass similarity between learned QPM features to that of genomic features.

## 2. Results

### 2.1. Bacterial genome assembly and bioinformatic analyses

Data from whole-genome alignment indicate an average intraspecies nucleotide similarity of 97.4%, 98.3%, 99.2%, and 93.7% between isolates belonging to *E. coli, A. baumannii, K. pneumoniae*, and *S. aureus* species (Fig 1). As expected, the only interspecies similarity was between *E. coli* and *K. pneumoniae* isolates (ca. 84%). Also, there was no ANI similarity among any of the other species. Furthermore, phylogeny analyses showed that all isolates from those four species were clustered together (Supplementary Figure 1). So, although the number of isolates is not large (21 isolates), the selected isolates have a broad molecular level diversity. Genome assembly quality analysis showed comparable results between isolates except for E. coli_NCTC_13441 (Supplementary Table 1).

**Fig 1.**
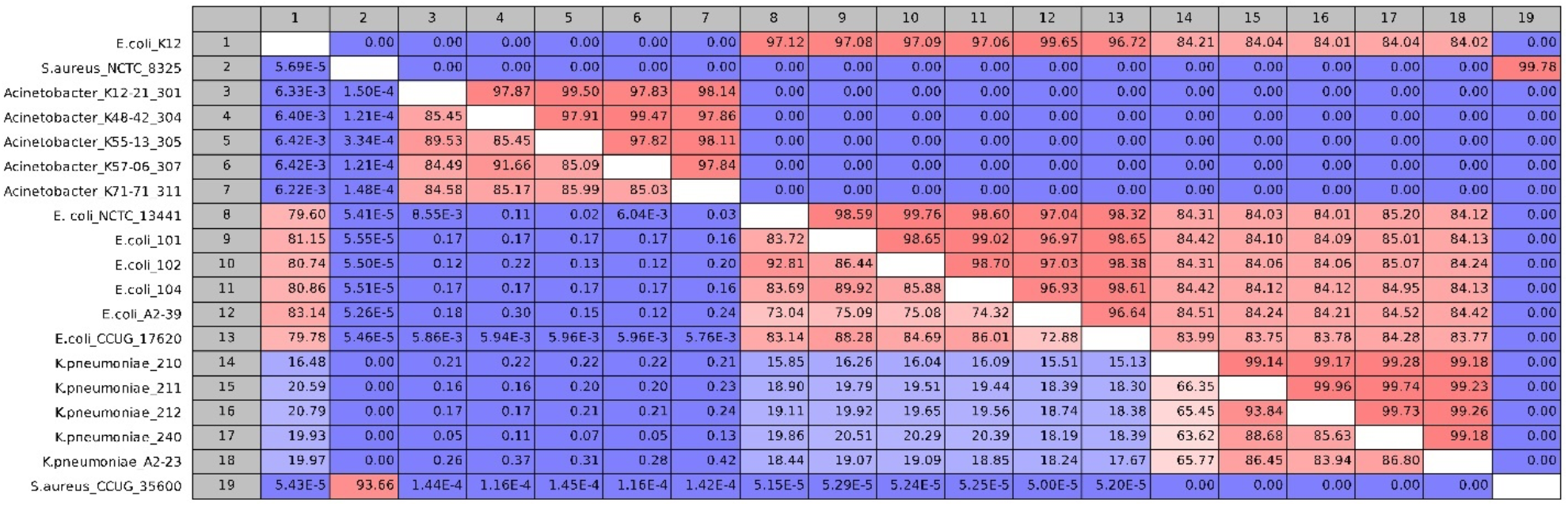
Whole-genome alignment for the 19 sequenced isolates. The upper and lower comparison gradient shows the percentage of nucleotide similarity and alignment percentage between genomes, respectively.

The five wild-type strains, including *E. coli* (CCUG17620), *E. coli* K-12, *Bacillus subtilis* (INN), and *A. baumannii* (INN), did not possess any antibiotic resistance genes (ARGs). In contrast, the *S. aureus* NCTC 8325 genome contained genome encoded *fosB* gene. A wide variety of ARGs conferring resistance towards penicillin, 3^rd^ generation cephalosporins, carbapenems, tetracyclines, and aminoglycosides was identified in the 16 resistant strains (Table 1). All the sequenced isolates possessed single or multiple plasmids except *A. baumannii* K48-42 and *E. coli* CCUG 17620. Putative plasmids in *E. coli* and *K. pneumoniae* isolates were previously validated using a plasmid-specific assembly approach [35]. In *A. baumannii* K48-48 and *E. coli* CCUG 17620, ARGs were chromosomally encoded. In methicillin-resistant *S. aureus* CCUG35600, the *mecA* gene was chromosomally encoded (Table 1).

**Table 1.**
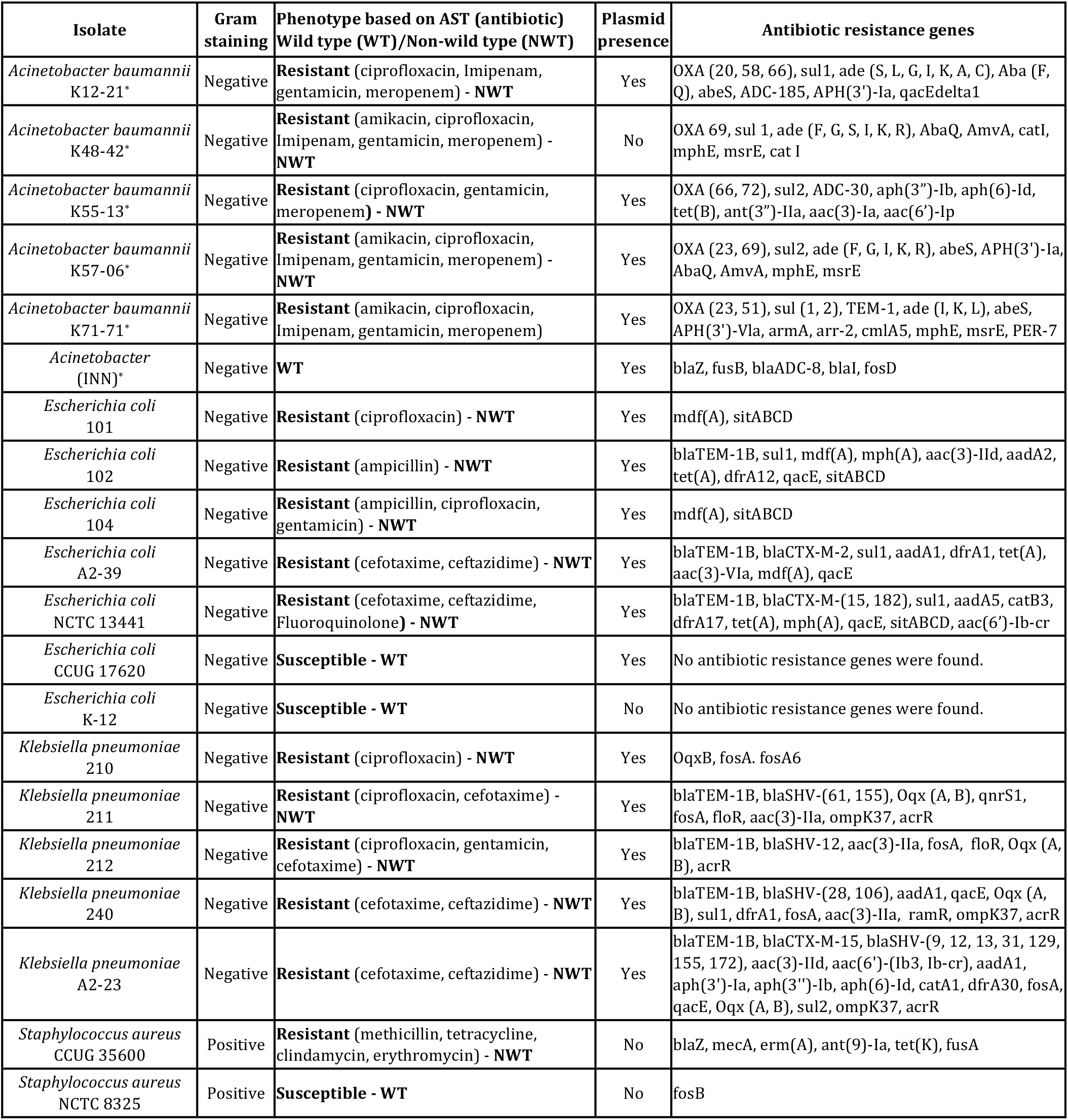
Overview of bacterial strains’ phenotype and genomic background. Phenotype and genotype information from antibiotic susceptibility testing (AST) and whole genome sequencing, respectively. ^*^*Acinetobacter has intrinsic resistance to several antibiotics*

### 2.2. Quantitative phase imaging of bacteria samples

To perform quantitative phase microscopy of bacterial samples, the sample is placed under the QPM setup shown in Fig. 8 for interferometric recording. The details of the bacteria sample preparation scheme, quantitative phase recovery, and defocus correction steps are provided in the Material and Methods section. The sample is placed on an XY motorized translation stage to acquire phase images of multiple FOVs using a high-resolution water immersion objective lens 60×/1.2NA. For each bacterial class, more than 50 different FOVs are acquired to generate large data sets. The numbers of bacteria present in one FOV were approximately equal to or greater than 300. This way, more than 15000 bacteria phase images of each class are generated for deep learning training. The recovered phase maps of different types of 21 strains of the bacterial samples are exhibited in Fig. 2.

**Fig. 2.**
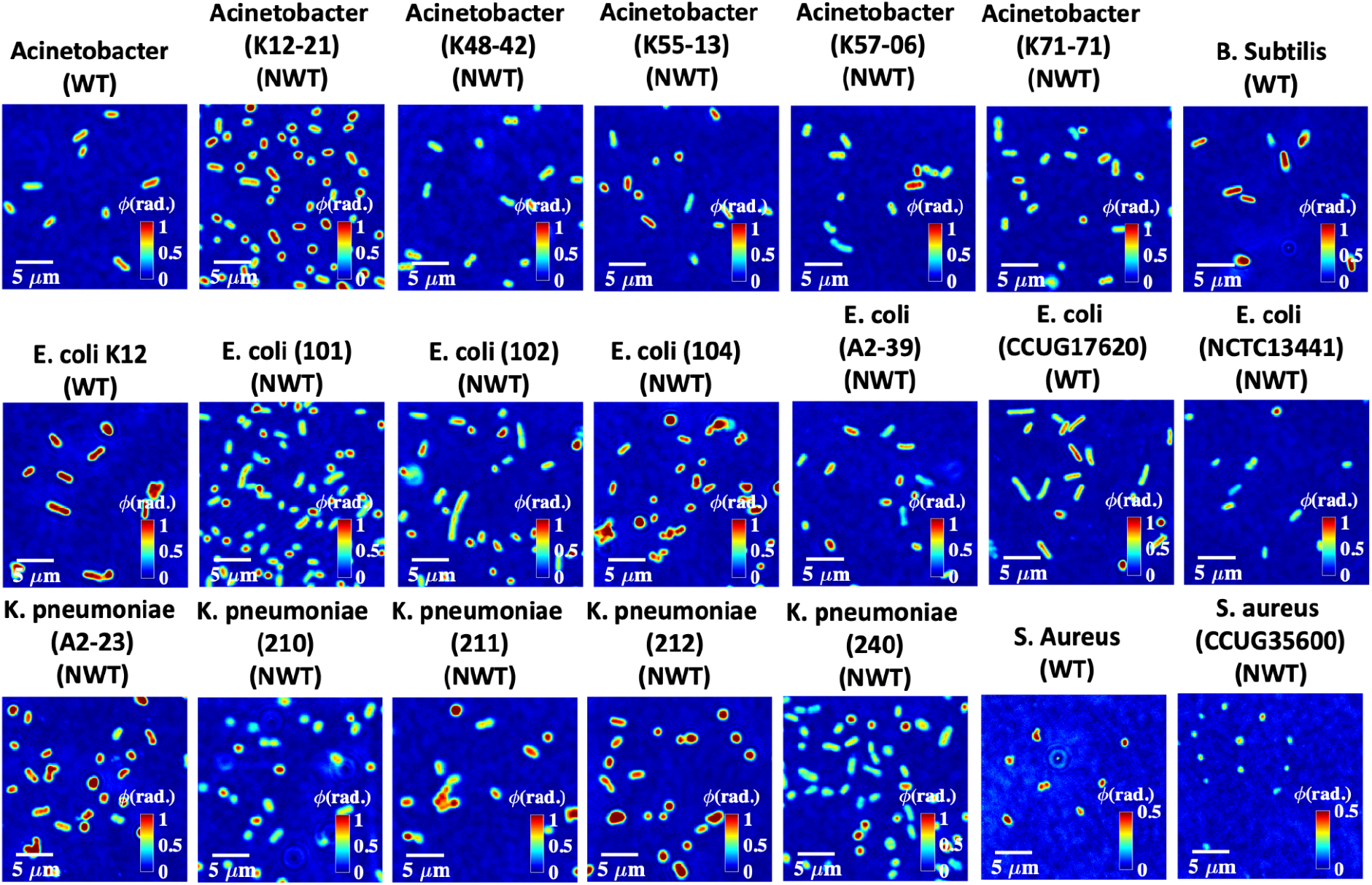
Quantitative phase maps of 21 different bacteria strains. The color bars for the bacteria samples are from 0 to 1 rad. Note that the color bars for *S. Aureus* strains are from 0 to 0.5 rad.

### 2.3. Defocus correction and segmentation at a single bacteria level

It is demonstrated in the Material and Methods section that sample defocus significantly affects the recovered phase profiles of the bacteria samples; therefore, it needs to be corrected before using bacteria phase images for the training of deep learning networks. In the Material and Methods section, the concept of defocus correction is demonstrated on a wide FOV. However, there might be different amounts of defocus in different regions of the recovered complex field over the entire FOV of the QPM system due to the presence of higher-order aberrations. The other source of error could be the slight defocus in the raw interferometric data itself due to the inability of the user to judge the best focus during acquisition correctly. Therefore, it is necessary to perform defocus correction at a single bacteria level to avoid any chances of error in the recovered phase maps.

Figure 3 illustrates the steps of the input phase data generation of the bacteria samples for deep learning network training. Firstly, the number of bacteria present in the entire FOV recovered phase maps are counted. Next, the positions of all the bacteria present in the entire FOV are stored and utilized to crop them. The sizes of the cropped images are not kept constant. It is decided by measuring the number of pixels covered by the bacteria cells along both rows and columns using the MATLAB *regionprops* command. The crop area is then decided by increasing 20 pixels along all four sides from the extreme coordinates of the bacteria. This helps to avoid any unwanted cropping of the bacteria region. Before cropping the bacteria phase images, the selection of the coordinates corresponding to each bacterium is made in a way such that the cropped images are of square shape by keeping the number of rows and columns equal. All the bacteria are kept at the center of the cropped phase image for further processing steps, as depicted in Fig. 3A.

**Fig. 3.**
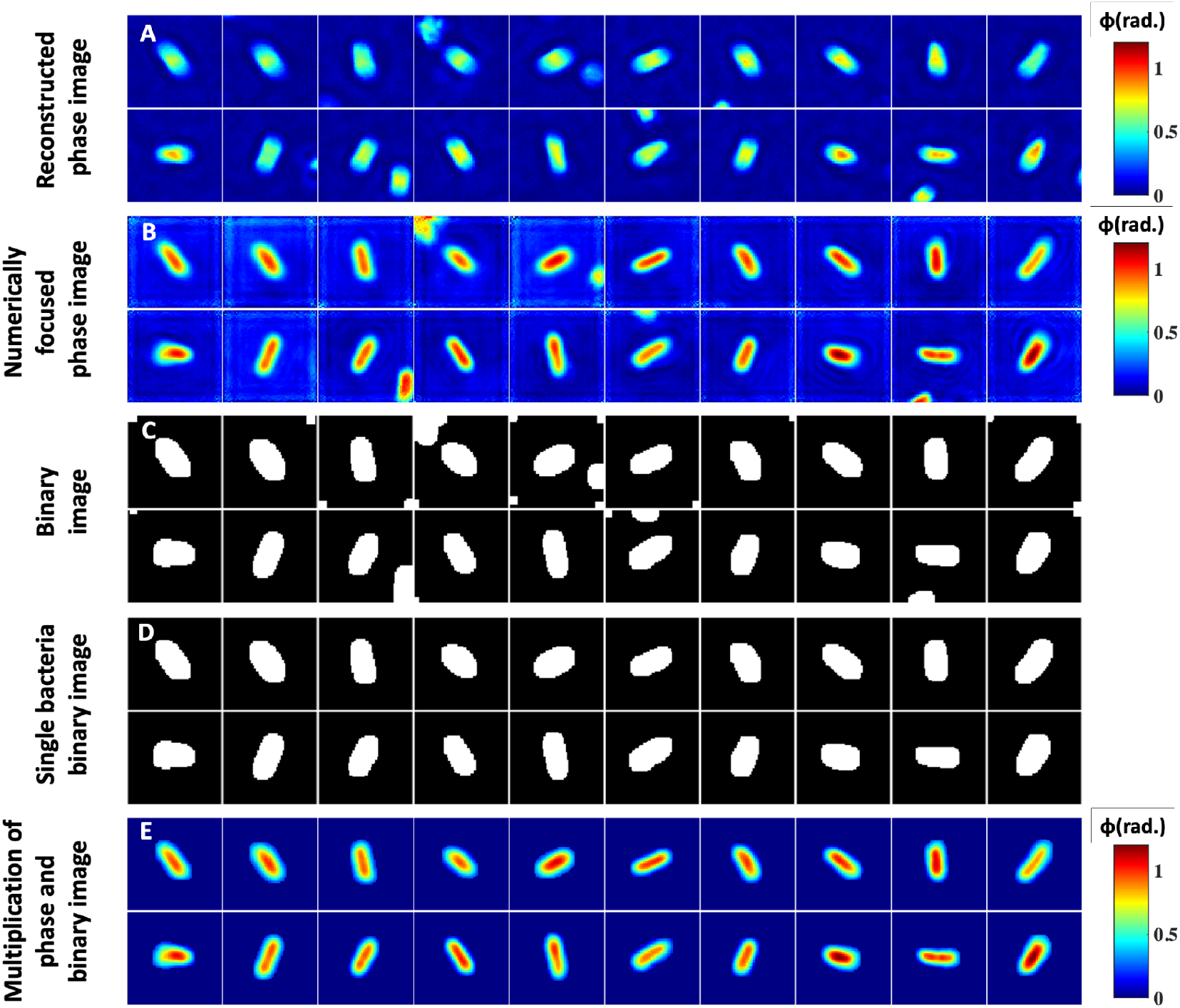
Segmentation and postprocessing pipeline for individual bacteria images. (A) Reconstructed phase maps of the individual bacteria cells. (B, C) Numerically focused phase images and corresponding binary maps. Some phase images have multiple bacteria at the edges, which need to be removed. (D) Represents a single bacteria binary mask. (E) Multiplication of the numerically focused phase maps and single bacteria binary masks.

Next, the amount of defocus in each cropped bacteria complex field is measured for the correction of defocus from the recovered phase images, if any. The numerically defocus corrected phase images of some of the bacteria samples are presented in Fig. 3B. The amount of defocus is found to be slightly different in all cropped bacteria phase images. It can be clearly visualized that defocus in the raw interferometric data significantly affected the recovered phase maps of the bacteria cells. Further, numerically focused phase images are utilized to generate corresponding binary images, as illustrated in Fig. 3C. To generate the binary images, the thresholding of the phase images is done by measuring the system’s peak to valley phase sensitivity. The threshold phase value is approximately 1.2-1.4 times the peak-to-valley phase error of the system for the bacteria segmentation. This threshold value is found to be an optimum value for the segmentation as the maximum phase values of most of the bacteria are approximately greater than 3-4 times the peak-to-valley phase error. The segmented areas of an image are further dilated by 2 pixels along all sides to remove any chances of unwanted bacteria phase map cropping. Some of the binary images have more than one isolated bacterium. The binary images are then improved by removing the regions of other bacteria using MATLAB, as shown in Fig. 3D. The generated binary image is then multiplied by the recovered phase image to remove the background regions from the phase images, as illustrated in Fig. 3E.

## 3. Deep learning for the classification of bacteria samples at colony and bacteria level

### 3.1. Data Preprocessing and model architecture

After segmenting bacteria to isolate multiple instances (Fig. 4(A2)), it was observed that there are still images with multiple bacteria co-located together, appearing as a single object. To exclude such images, we employ a heuristic-based approach that considers the area of the object and its convex hull. As illustrated in Fig. 4 (A and B), we obtain the object contour and the convex hull. By considering the area of these two polygons, we derive a normalized score as,

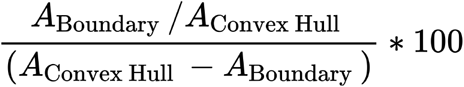

Based on this score and a selected threshold for the dataset, we remove images that have co-located bacteria Fig.4 (A and B). After this filtering process, 472,795 images were selected, resized to 224 × 224, and then split according to the ratios 80%, 10%, and 10% for training, validation, and testing, respectively. Stratified sampling ensures that the identical class distributions are preserved in these 3 data splits.

**Fig. 4.**
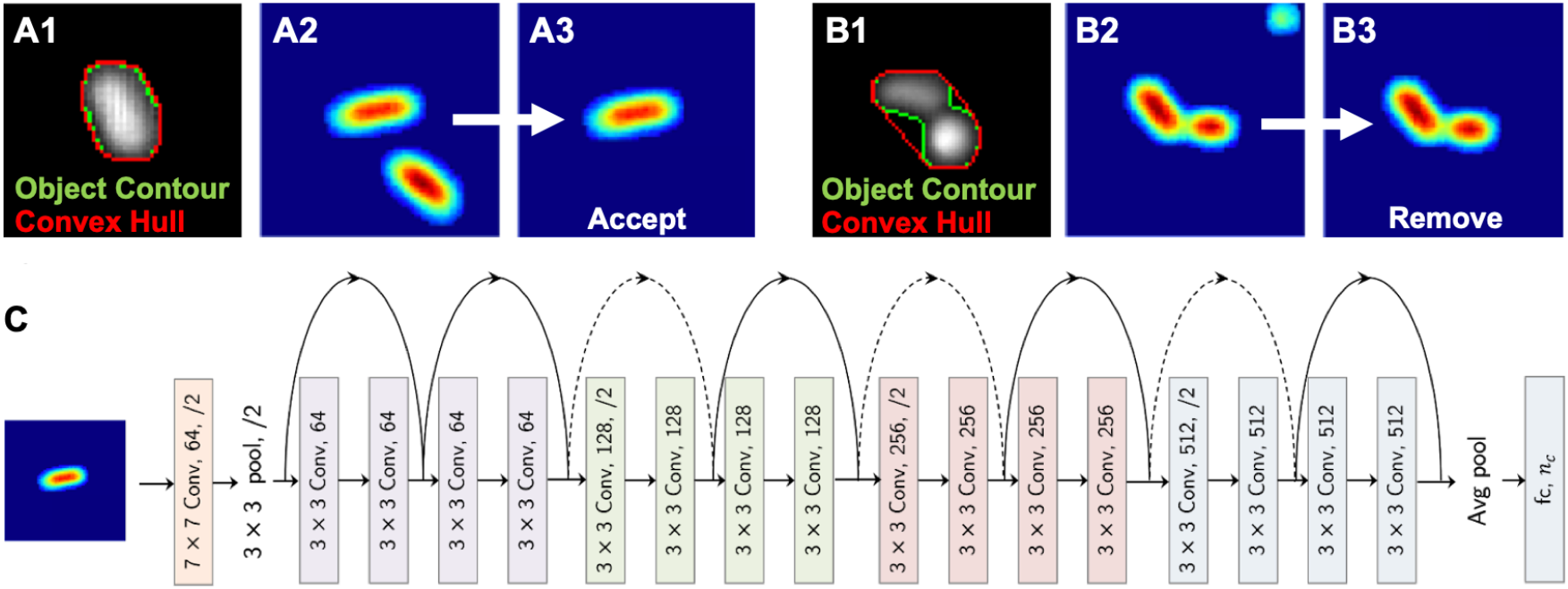
Deep neural network pipeline. **(A, B)** A heuristic for detecting co-located bacteria. **(C)** Neural network architecture.

We employ the deep residual learning framework introduced by He *et al*. ^(11)^, which can be used to overcome the performance degradation problem that arises when training deeper networks. The 18-layer residual network (Fig. 4(C)), which was pretrained on the imagenet dataset, is used as the initialization for the weights and is fine-tuned on our dataset of bacteria images. The model weights are optimized using the stochastic gradient descent optimizer with a learning rate of 0.01 and a momentum factor of 0.9. The overall model architecture is shown in Fig. 4. Based on our classification requirement (binary vs. multiclass classification), the number of output neurons (n_c_) are defined. For both antibiotic resistance prediction and gram stain classification, n_c_ = 2, while for species-level classification and strain classification, the number of output neurons is 5 and 21 based on the number of output classes, respectively.

### 3.2. Experiments & Evaluation Criteria

The model architecture described in Section 3.1 is adopted and trained to measure the performance on classification tasks: 1) Antibiotic resistant/sensitive prediction, 2) Gram stain classification, 3) Species-level classification, and 4) Strain classification. A separate classifier was trained for 8 epochs for each task. The trained model measures the performance of bacteria samples from a blind test set. We varied the number of bacteria images in the sample as N = 1, 3, 7, 15, 31, and 63 (also 127 for the strain level classifier) to gauge the minimum number of bacteria that should be imaged for each classification task. Each image of bacteria was classified independently using the classifier, and individual predictions on these images were aggregated to obtain the prediction per sample (at each N).

We first analyze the antibiotic resistant/sensitive and gram-stain prediction, which are binary classification tasks. Fig. 5A shows the results of antibiotic resistance prediction. The positive (resistant class) prediction achieved an accuracy of 100% (i.e., the ratio between correctly predicted tests out of total tests from the positive class) at N=15, while the non-resistant prediction accuracy was 96.45% at N=63. In the gram stain prediction task (Fig. 5B), the gram-negative class achieved 100% accuracy at N=7, while the gram-positive prediction was 83.43% at N=31.

**Fig. 5.**
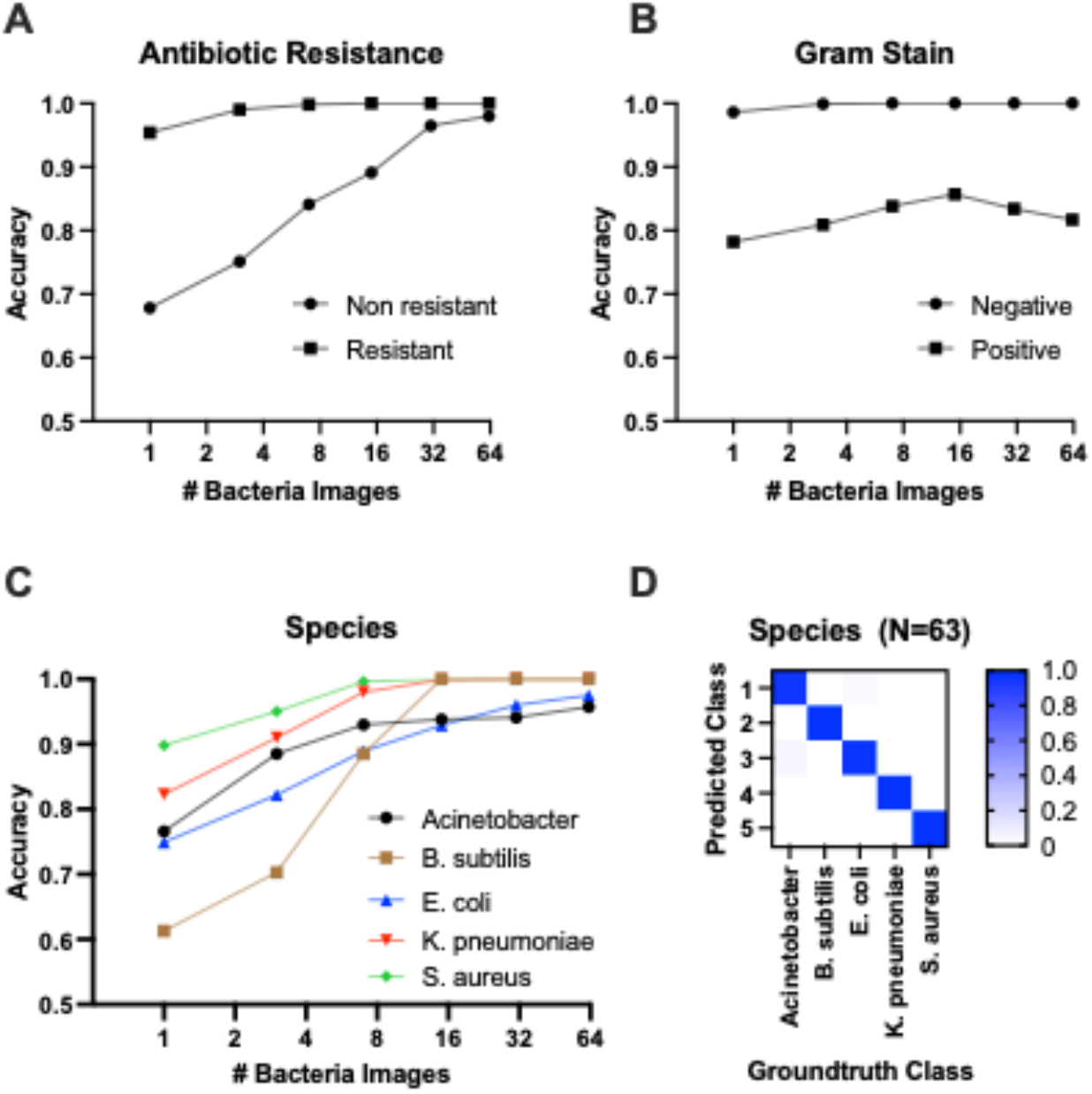
Classification results against the number of bacteria images. **(A)** Classification accuracies for antibiotic resistance. **(B)** Classification accuracies for gram +/-ve. **(C)** Species-level classification accuracy. **(D)** Confusion matrix for species-level classification with 63 bacteria images in the batch.

Second, we present the results for species-level classification (Fig. 5C). The model predicted *B. subtilis, K. pneumoniae*, and *S. aureus* classes with 100% accuracy at N=31. For *A. baumannii*, the accuracy was 0.957 at N=63. For *E. coli*, the accuracy was 0.975 at N=63. The confusion matrix for species-level classification at N=63 is shown in Fig. 5D.

Next, Fig. 6 (A and B) shows the strain level classification results. The model was 100% accurate in predicting 18 out of the 21 strains at N=31 (Fig. 6A). Considering challenging strains, the classification accuracy increased with increasing the N for E. coli 102 and E. coli A2-39. At N=127, the accuracy of *E. coli 102* and *E. coli A2-39* reached 100% and 73%, respectively. However, the classification accuracy of *A. baumannii* K55-13 remained consistently below 35% with increasing N. Fig. 6B shows the confusion matrix for strain level classification at N=63. As seen at N=63, a little under 30% of *E. coli* 102 samples were misclassified as *E. coli* CCUG17620. Also, ca. 35% of *E. coli* A2-39 samples were misclassified as *E. coli* NCTC13441. Finally, almost 70% of *A. baumannii* K55-13 samples were misclassified as *E. coli* A2-39.

**Fig. 6.**
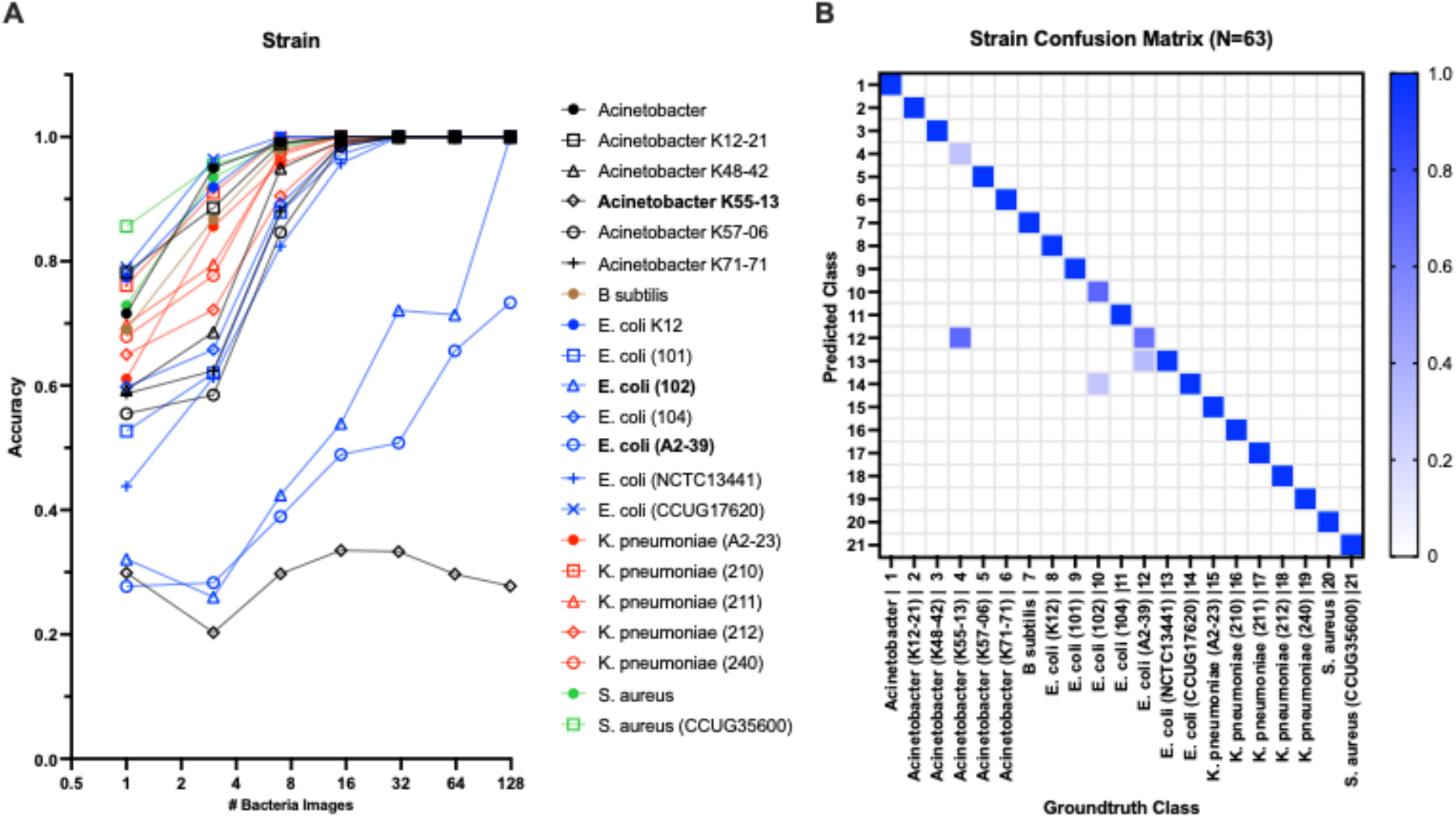
**(A)** Strain level classification accuracy against the number of bacterial QPM images **(B)** Confusion matrix for strain-level classification with 63 bacterial QPM images in the batch. The WGS data provide the ground truth class information.

Finally, we investigated the probability of a QPM image from each strain being classified as one of the 21 strains as a similarity measure of QPM features. For each test image from a particular class, the probability of prediction was extracted and averaged over all images from that class. Fig. 7B shows the QPM similarity matrix. Fig. 7A illustrates the equivalent similarity measure from the genomic analysis; here, the average nucleotide identity (ANI) was used. Shown in red boxes are the similarities between the strains from each species. As expected, ANI was more than 90% similar within the species. Some strains from *Acinetobacter* (K48-42, K55-13, K57-06, and K71-71), *E. coli* (101, 102, 104, and CCUG17620), *K. pneumoniae* (A2-23, 211, and 212), and *S. aureus* (NCTC 8325 and CCUG35600) showed some QPM similarity. Nevertheless, no distinct patterns were observed between the ANI and QPM similarities.

**Fig. 7.**
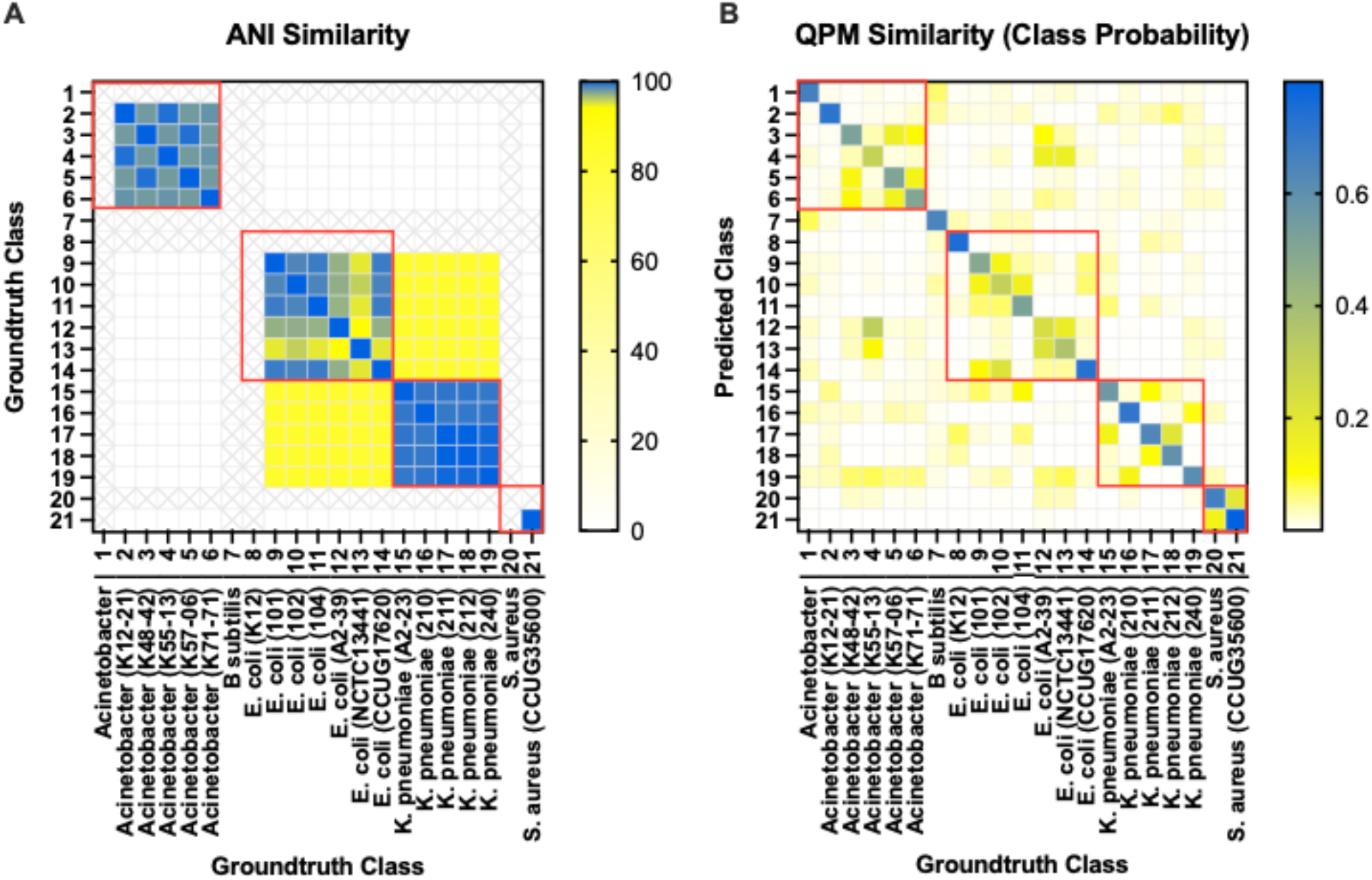
**(A)** Average nucleotide identity (ANI) based similarity between bacteria classes. Classes representing isolates that weren’t sequenced are shown in ‘X’s. **(B)** The equivalent QPM-based similarity between bacteria classes. Here the probability of strain level class predictions (averaged over all test instances for each class) was used as the similarity measure.

**Fig. 8.**
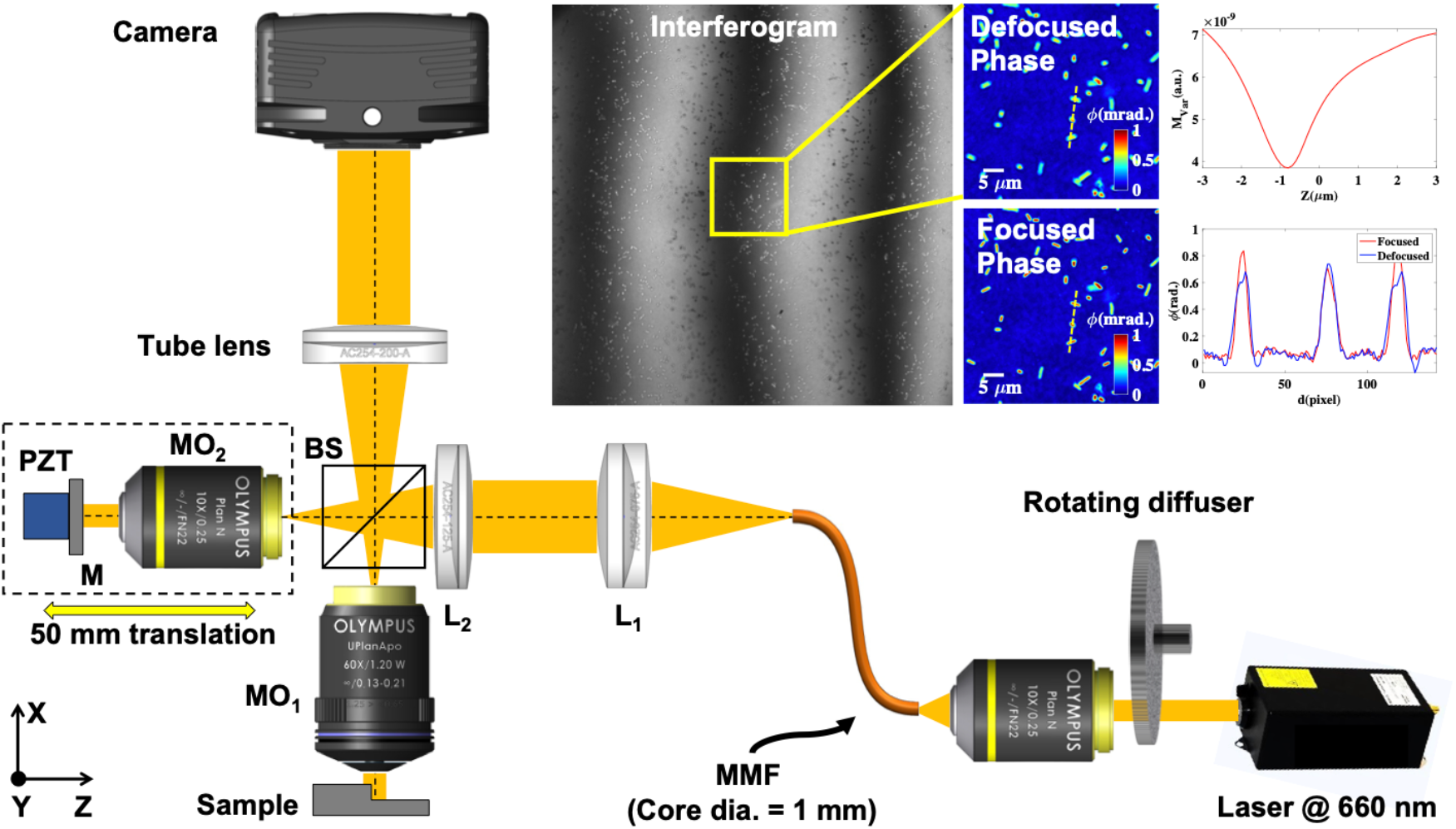
Optical schematic diagram of Linnik interferometer-based QPM setup. MO_1-2_: Microscope objectives; BS_1-2_: Beam splitters; L_1-2_: Achromatic doublet lenses; MMF: Multi-mode fiber; M: Mirror and PZT: Piezo electric transducer. The inset represents the interferogram, reconstructed defocused phase map, numerically focused phase map, and amplitude variance plot of the numerically propagated optical field as a function of the propagation distance. The line profiles of the bacteria phase map before and after focus correction along the yellow dotted lines.

## 4. Discussion and conclusion

A significant challenge for clinicians managing infection is to precisely identify those patients who should receive antibiotics (and which one) and those who should not. This is due to physicians not being able to diagnose patients accurately in real-time, leading to either prescription of antibiotics for viral infections or a prescription of broad-spectrum drugs that should ideally be kept in reserve. The current state-of-the-art diagnostic of infection is mainly based on biochemical analysis of the culture of the clinical material. This leads to a detection time of 48-96 hours or even longer.

In this study, we imaged isolates from 21 unique bacterial strains from five different species, including four WHO priority pathogens (*E. coli, A. baumannii, K. pneumoniae*, and *S. aureus*) using a PTLS based high-sensitive quantitative phase microscope (QPM). PTLS brings many advantages, such as high-sensitivity, high spatial and temporal resolution, and the use of a high-resolution objective lens in QPM. First, many sources of errors, such as higher-order phase aberration and defocusing effect on the phase maps of the bacteria samples, are fixed. These are crucial steps and need to be adopted before using the QPM images of bacteria samples in deep learning, which may mislead its classification outcome. Using a deep neural network, we then classified the QPM images of the individual bacterium at strain and species levels and according to their antibacterial resistance and gram-staining. WGS data (from Illumina and MinION sequencing platforms) and AST data were used to generate the ground truth (strain, species, and resistance profile) to train the DNN.

Our results showed high classification accuracy for antibiotic resistant/sensitive, gram staining, and species identification. The network could also classify 19 out of the 21 strains when at least 128 bacteria images were analyzed. One of the misclassifications was of *E. coli* A2-39 samples which were misclassified as *E. coli* NCTC13441 and the other were *A. baumannii* K55-13 samples which were misclassified as *E. coli* A2-39. Amongst these the misclassification of *A. baumannii* as *E. coli* is clinically misleading as these are two different species, which would lead to prescription of the incorrect antibiotic.

These results show the potential of the current approach as bacteria cell culturing would not be needed, which is a time-consuming process, for the identification of different bacteria samples at strain and species level. It is observed that out of two strains that are either weekly classified or not classified, for one strain, the results may be improved by analyzing more bacteria; nevertheless, for the other strains, the network was incapable of correct classification. The two misclassified classes on our training set were due to their similarity in QPM images, which demonstrates the limitations of deep-learning-based methods with limited image information.

Our results suggest that differential morphological features from different bacteria strains can be measured using our approach, and they can be used in downstream machine learning pipelines for detection. These findings demonstrate the utility of using QPM and DNN to identify a range of WHO priority pathogens and provide relevant information about microbiological samples, which WGS or conventional microbiological methods can later verify. Our results establish the potential of QPM as a rapid first-stage analytical tool that can complement WGS for phenotype prediction and resistome analysis. Such a workflow can be hugely impactful in handling and preventing the spread of AMR and could lead to future use in clinical microbiology.

To the best of our knowledge, this is the first study that has used a combination of enabling technologies using genomics, QPM, and deep learning to rapidly detect pathogens and their resistance profile. Moreover, in addition to the species-level classification, which is frequently studied in the literature, we have analyzed the classification performance under multiple levels of classification tasks, including presence or absence of resistance, gram staining, and strain-level classification, which has been followed up by WGS data and AST for ground truth information to validate the results. Taken together, all this information is of high clinical importance and can be utilized as a heuristic approach compared to state-of-the-art solutions.

## Material and Methods

### Bacterial strains and phenotypic assessment

The wild-type/sensitive and phenotypically resistant/non-wild-type strains of Gram-negative *Acinetobacter baumannii* (n=6), *Escherichia coli* (n=7), *Klebsiella pneumoniae* (n=5), and Gram-positive *Staphylococcus aureus* (n=2), *Bacillus subtilis* (n=1) were used for this study Phenotypic minimum inhibitory concentration (MIC) information on the *E. coli* and *K. pneumoniae* strains was taken from the reference strain collection or has been previously published [36]. MIC values of Acinetobacter strains were as described in [37] and for *S. aureus* strains (https://www.ccug.se/strain?id=35600 and https://bacdive.dsmz.de/strain/14451). Isolates were classified as susceptible and resistant according to the European Committee on Antimicrobial Susceptibility Testing (EUCAST) Breakpoints v 12.0 (December 2021).

### Sequencing and bacterial genome assembly

The Acinetobacter isolates libraries were prepared with the Nextera XT DNA Library preparation kit from Illumina (Cat. No.: FC-131-1096) and were sequenced with the NextSeq 550 instrument (Illumina, USA), using PE mode and a mid-output flow cell (150 cycles). The WGS data for E. coli CCUG 17620, E. coli NCTC 13441, E. coli K-12, and S. aureus NCTC 8325 strains were downloaded from NCBI (BioSample: SAMN0993043, SAMEA2432036, SAMN02604091, SAMN02604235). Sequencing data for the remaining samples were taken from our previously published work [36, 38]. The Illumina reads were first quality checked using FastQC (v 0.11.8 for Linux) [39]. Adapters were removed, and low-quality reads (Phred <25) were filtered out using Trimmomatic [40] with default parameters. For MinION reads, adapter and barcode trimming were performed using Porechop (v0.2.4 for Linux) with default settings [41]. Long and high-quality reads were collected using Filtlong (v0.2.0 for Linux) with default parameters [42].

According to our previous data, Unicycler and Flye assemblers outperformed other short and long-read assemblers [43]. Therefore, in this study, genome assembly was performed using Unicycler (v0.4.9) and Flye (v2.8.2) for short and long reads, respectively [44, 45]. The assemblies were quality checked using QUAST (v4.6.0 for Linux) in addition to BUSCO [46-48] and annotated using Prokka (v1.14.5 for Linux) [49]. The whole-genome alignment was performed, and the phylogeny tree was constructed using the CLC Genomics Workbench version 21.0.3 (QIAGEN) [50]. Using the whole-genome alignment toolbox in CLC, the average nucleotide identity (ANI) and alignment percentage (AP) were obtained for paired comparison between samples. Later the identity score matrix from ANI was used for phylogenic tree construction using the neighbor-joining approach.

### Identification of plasmids and antibacterial resistance genes

FASTA files of *De novo* assembled genomes were used to identify plasmids and antimicrobial resistance genes (AMR). To identify plasmids, the PlasmidFinder online tool (software version: 2.0.1, database version: 2020-07-13) [51], with a minimum identity of 95% and coverage of 60%, was utilized for all isolates, except *A. baumannii*, where the plasmids were identified using BLAST search against the PLSDB database [52].

For *E. coli, K. pneumoniae*, and *S. aureus* isolates, antibacterial resistance genes associated with mobile elements were identified using Resfinder online tool (v 4.1, software version: 2020-10-21, database version: 2020-12-01) [53]. The PointFinder online tool (software version: 2020-10-21, database version: 2019-07-02) [54] was used to predict antibacterial resistance genes associated with chromosomal point mutation. The AMR genes in *A. baumannii* isolates were identified using the Resistance Gene Identifier tool in CARD v3.1.4, update version 2021-10-05 [55]. Only hits flagged as perfect were considered true resistance genes.

### Culture and fixation of bacterial cells for microscopy

The bacterial strains were cultured from frozen aliquots (−80 °C) on blood agar plates, and the cultures were used after 20 hours of incubation at 37 °C. After overnight culture, the OD600 nm was measured for each strain, and the cells were collected and re-suspended in 1 ml PBS to reduce the presence of agar components within the samples. The cells were fixed with a ratio of 1:2 2% paraformaldehyde in phosphate sodium buffer (PFA in PBS, pH 7.15) and incubated at room temperature for 1 hour. The cells were washed twice and re-suspended in 500 μl PBS and transferred to sterile tubes for shipping for QPM analysis.

### Bacteria immobilization for quantitative phase imaging

Bacteria immobilization is a crucial step for their high-resolution quantitative phase imaging. High-resolution phase recovery requires multiple phase-shifted interferograms; therefore, the bacteria cells should not move during the data acquisition. Due to their small size, bacteria have Brownian motion and quickly change their position and orientation. Therefore, bacteria immobilization is needed so that reliable data can be recorded.

As Linnik interference microscopy configuration works in the reflection mode, therefore, the bacteria samples are prepared on a reflecting substrate (Si-wafer). The Si-wafer substrate is first incubated with 0.01% poly-L-lysine (PLL) solution for 15 - 20 min. The excessive PLL solution is removed, and the substrate is washed thoroughly with phosphate-buffered saline (PBS). It formed a thin layer of PLL on the substrate and made it positively charged. The polydimethylsiloxane (PDMS) chamber of 10 mm × 10 mm size with 150 μm thickness is placed on the Si substrate. The bacteria samples were seeded into the PDMS chamber and left for approximately 30 min for their incubation. The electrostatic attraction between the negatively charged bacteria and positively charged PLL solution helped them to adhere to the Si substrate [56].

The adherence of some species/strains of the bacteria to the Si substrate was observed to be a little challenging. Different strains of *E. coli* and *K. pneumoniae* were attached straightforwardly on Si substrate after following the aforementioned protocol. In the case of *S. aureus, A. baumannii*, and *Bacillus subtilis*, the attachment was challenging. Several factors, such as the chemical constituents of liquid, morphological features of bacteria, and the incubation time, could affect the immobilization process [56]. For these classes, the Si substrate is seeded with concentrated sample volume and incubated for an extended period, around 1 hour. The sample is gently washed off with PBS to remove mobile bacteria cells. The substrate is then left only with the bacteria cells attached to the substrate. PBS is added to the sample and sealed with # 1.5 cover glass from the top, which enabled the use of a water immersion objective lens for imaging and avoided any air current in the specimen.

### Experimental scheme of quantitative phase microscopy

The experimental scheme of the QPM system is illustrated in Fig. 8, which is utilized to acquire phase-shifted interferometric data of the bacterial samples. The working principle of the QPM system is based on a high-resolution Linnik interference microscopy configuration. In the QPM system, a temporally high and spatially low coherent light source, also called pseudo-thermal light source (PTLS), is utilized due to various advantages over conventional light sources like a laser, halogen lamps, and LEDs light sources. PTLS enables high-resolution phase recovery of biological specimens with high spatial phase sensitivity. PTLS is generated when a high temporal coherent laser beam is passed through a rotating diffuser. The output of the rotating diffuser acts as a temporally high and spatially low coherent light source and enables speckle noise and a coherent noise-free phase recovery [23].

In our experimental scheme, a laser light beam at 660 nm wavelength (Cobolt Flamenco) was passed through a rotating diffuser, and the output of the diffuser was coupled into a multi-mode fiber (MMF) of a core diameter of 1 mm using an objective lens 10×/0.25NA. The output of MMF was first nearly collimated using lens L1 and then focused using lens L_2_ at the back focal plane of the object arm objective lens (MO_1_: 60×/1.2NA) to achieve uniform illumination at the specimen. The focused light beam was split into the object and the reference beam using beam splitter BS. The reference beam was passed through a reference arm objective lens (MO2: 10×/0.25NA) and reflected from the mirror. The light beam is again collected by the same objective lens and called a reference beam. The objective lens in another arm captured the bacteria sample information, which was recombined with the reference beam at BS and projected at the camera sensor using a tube lens (TL) to form the interferograms. The interferograms were further captured by employing a scientific Hamamatsu CMOS camera. The reference mirror was attached to a nanometric precision piezo stage to introduce a phase shift between the interfering beams to acquire multiple phases shifted interferograms of the specimen for phase recovery. A homemade LabVIEW software program was written and utilized for the required phase stepping in the interferograms and their acquisition using a camera. The reference mirror was also attached to a kinematic mirror mount to control the angle between the object and the reference beam. The total acquisition time of 5-6 phase-shifted interferograms was approximately 600 ms, which can be improved using a high-end computer.

### Quantitative phase microscopy of bacteria and defocus correction

For phase recovery of the sample, multiple phases shifted frames are recorded to achieve diffraction-limited phase recovery. One of the phase-shifted interferograms is depicted in Fig. 9A. It can be seen that interferograms suffer from higher-order phase aberration and would influence the recovered phase maps. Multiple phase-shifted interferograms are utilized to recover the complex field information of the bacteria sample using the principal component analysis (PCA) algorithm [57]. Full field of view (FOV) amplitude and phase part of the recovered complex field is illustrated in Figs. 9 (B and D). The zoomed views of the regions marked with yellow color boxes in the amplitude and phase parts are depicted in Figs. 9 (C and E), respectively. It can be clearly visualized that the recovered phase map looks defocused, which affected the shape of the bacteria significantly. Therefore, it becomes necessary to perform numerical defocus correction steps to achieve faithfully reconstructed phase maps. The defocus correction algorithm steps are given in the Supplementary information. Fig. 9F presents the numerically focused phase image. After following the numerical defocus correction steps, the bacteria phase image looks sharply focused. The amount of defocus is found to be equal to 1 µm by observing the sharpness curve. In addition, defocusing of the bacteria significantly affected the morphological parameters such as area, maximum phase value, volume, etc. Thus, this is a crucial step for the accurate classification of different classes of bacteria using deep learning. Otherwise, there would be a possibility of misclassification of different classes of bacteria samples.

**Fig. 9.**
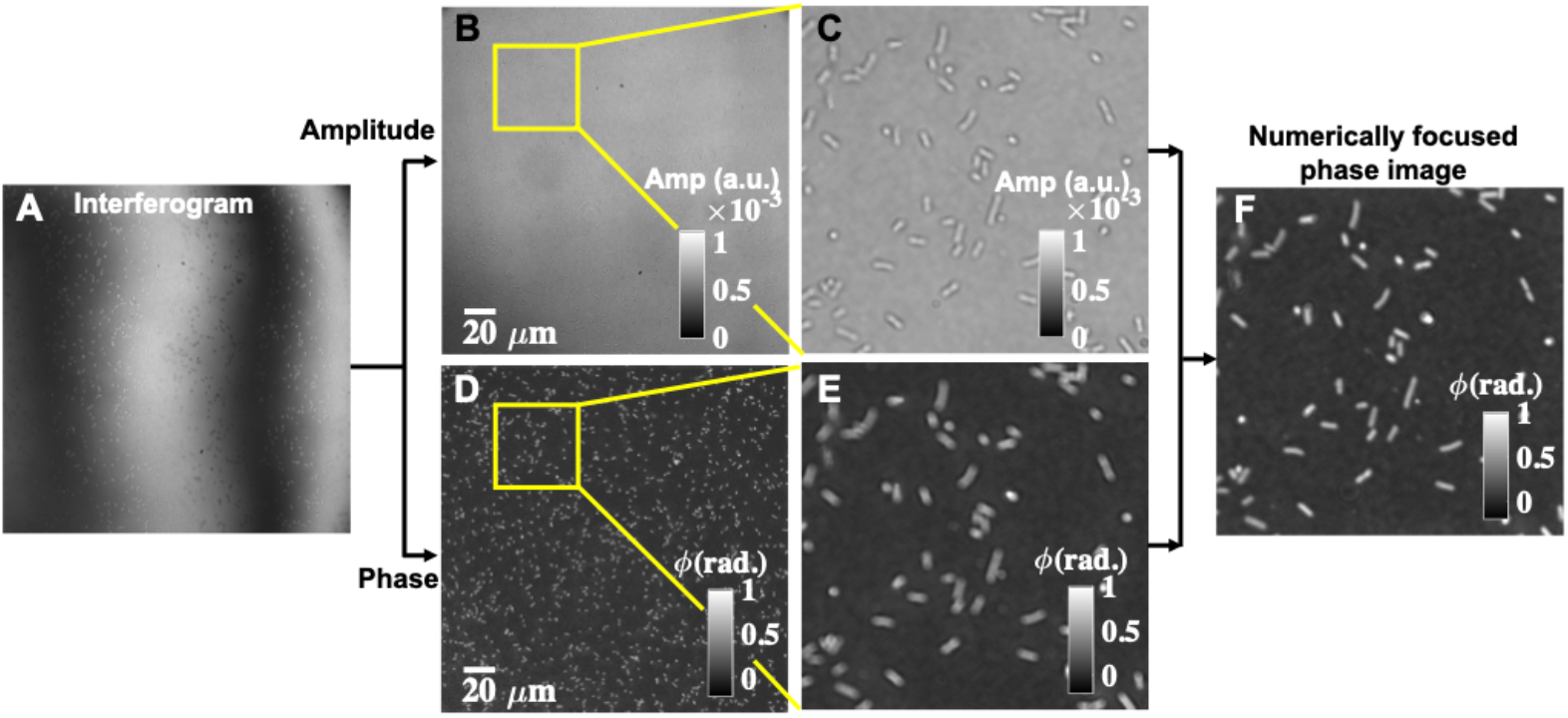
Reconstruction of numerically focused phase images using interferometric measurements from the QPM microscope. (A) One of the phase-shifted interferograms. (B, C) Corresponding amplitude and phase map of the bacteria sample. (C, E) Zoomed views of the amplitude and the phase images of the regions marked with yellow color boxes. (F) Numerically focused phase map.

## Notes

### Competing Interest Statement

The authors have declared no competing interest.

### Summary of Updates

Updated Table 1 and methodology.

